# Correlation between cancer stem cells, inflammation and malignant transformation in a DEN-induced model of hepatic carcinogenesis

**DOI:** 10.1101/2020.04.17.046391

**Authors:** Chun-Chieh Wu, Chien-Ju Lin, Kong-Kai Kuo, Wan-Tzu Chen, Chen-Guo Ker, Chee-Yin Chai, Hung-Pei Tsai, Sheau-Fang Yang

## Abstract

**Aims:** Chronic inflammation and cancer stem cells are known risk factors for tumorigenesis. The aetiology of hepatocellular carcinoma (HCC) involves a multistep pathological process characterised by chronic inflammation and hepatocyte damage but the correlation between HCC, inflammation and cancer stem cell remains unclear. In this study, we examined the role of hepatic progenitor cells in a mouse model of chemical-induced hepatocarcinogenesis to elucidate the relationship between inflammation, malignant transformation and cancer stem cells.

**Methods and results:** We used diethylnitrosamine (DEN) to induce liver tumour and scored for H&E and reticulin staining, and also immunohistochemistry staining for OV-6 expression and analysed the statistical correlation between each other. DEN progressively induced inflammation at 7 week 7 (40%, 2/5), week 27 (75%, 6/8), week 33 (62.5%, 5/8) and week 50 (100%, 12/12). DEN progressively induced malignant transformation at week 7 (0%, 0/5), week 27 (87.5%, 7/8), week 33 (100%, 8/8) and week 50 (100%, 12/12). Data obtained showed that DEN progressively induced high-levels of OV-6 expression at week 7 (20%, 1/5), week 27 (37.5%, 3/8), week 33 (50%, 4/8) and week 50 (100%, 12/12). DEN-induced inflammation, malignant transformation and high-level OV-6 expression in hamster liver as shown above and applying Spearman’s correlation to the data showed that expression of OV-6 was significantly correlated to inflammation (*p* = 0.001) and malignant transformation (*p* < 0.001)

**Conclusions:** There was a significant correlation between number of cancer stem cells, inflammation and malignant transformation in a DEN-induced model of hepatic carcinogenesis in the hamster.

## Introduction

Hepatocellular carcinoma (HCC) is a common malignancy that affects nearly one million people globally each year and treatment options are limited^1^. The aetiology of HCC involves a multistep pathological process characterised by chronic inflammation and hepatocyte damage^2^. The dedifferentiation of mature liver cells has been implicated in HCC^3^, based on which it was hypothesised that HCC arises from maturation arrest in liver stem cells and hepatic progenitor cells^4^. In light of this evidence, in this study we examined the role of hepatic progenitor cells in a mouse model of chemical-induced hepatocarcinogenesis.

Cancer stem cells are tumour cells with the ability to self-replicate and differentiate into solid tumours, including HCC^3^, and express known stem cell markers, such as CD133 in glioma, CD44 and CD24 in breast cancer and OV-6 in HCC^5–7^.

Oval cells are associated with the intrahepatic biliary system and are derived from hepatoblasts located near hepatic portals. These cells and their progeny have the ability to proliferate and differentiate into either biliary cells or hepatocytes. In the mature liver, oval cells induce the replication of hepatocytes and cholangiocytes^8^. Oval cells have been demonstrated to arise in rat hepatic intraportal regions after treatment with hepatocarcinogens or hepatotoxins^8, 9^. The cellular protein, OV-6, has been found to be a useful marker of rat oval cells, thought to be the progeny of hepatic stem cells^10, 11^. Intraperitoneal injection of mice with diethylnitrosoamine (DEN) was shown to induce tumours in the liver, gastrointestinal tract, skin, respiratory tract and haematopoietic cancer^12–14^. In this study, (DEN) was used as a carcinogen in an experimental hamster model to induce liver tumour and determine the relationship between malignant transformation, inflammation and the role of cancer stem cells in the pathogenesis.

## Materials and Methods

### Animals

Male hamsters (n = 40) were housed in plastic cages with soft bedding under a 12 hr reversed light-dark cycle (light cycle from 6 am to 6 pm; dark cycle from 6 pm to 6 am) and access to food and water *ad libitum*. All experimental procedures were approved by the Kaohsiung Institutional Animal Care and Use Committee.

### DEN-Induced Liver Cancer Model

The DEN-induced liver tumour model was established as described previously^12–14^. Hamsters were fed DEN (80 μg/g body weight/day). The animals were sacrificed at various timepoints in the study (weeks 7, 27, 33 and 50), and the liver was removed and fixed in 4% paraformaldehyde and embedded in paraffin.

### Histology analysis

Tissue sections (4 μm) were stained with haematoxylin and eosin (H&E) and by immunohistochemistry for various cellular markers. Tissues were stained for reticulin, which reflected the distribution of reticulin in the normal liver, where the sinusoids are lined by a rich reticulin framework; periportal reticulin is discrete without projections into the parenchyma. Each pathologist initially reviewed all slides and reached an agreement on the scores. Portal inflammation was scored from 0 to 3 where a score of 0 indicated no portal inflammation (Fig. 1A), score 1 indicated ≤2 portal inflammation (Fig. 1B), score 2 indicated >2 portal inflammation (Fig. 1C) and score 3 indicated portal inflammation with interstitial sinusoidal inflammation (Fig. 1D).

**Fig. 1.**
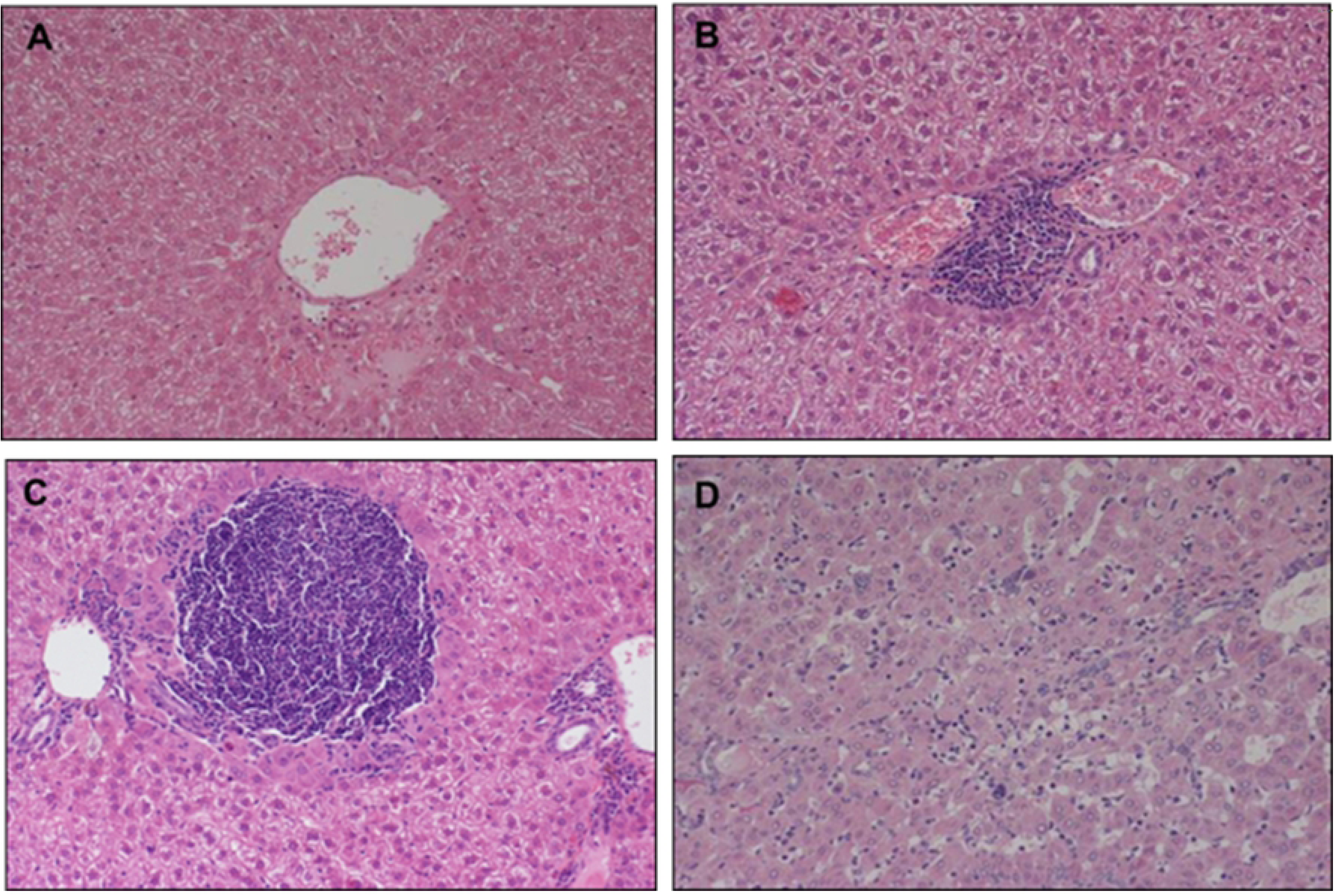
H&E staining of hamster liver: Portal inflammation was scored from 0 to 3 where a score of 0 indicated no portal inflammation (Fig. 1A), score 1 indicated ≤2 portal inflammation (Fig. 1B), score 2 indicated >2 portal inflammation (Fig. 1C) and score 3 indicated portal inflammation with interstitial sinusoidal inflammation (Fig. 1D).

### Reticulin Staining

Tissue sections measuring 4 µm obtained after deparaffinisation and rehydration from paraffin-embedded samples were rinsed in distilled water, and immersed in 1% potassium permanganate for 2 min followed by treatment as follows: 2.5% oxalic acid for 1 min, 2% iron alum for 1 min, Gomori’s solution for 3 min, 10% formalin for 2 min, gold chloride (1:500) for 3 min, 3% potassium metabisulfite for 1 min and 3% sodium thiosulfate for 1 min. Slide sections were rinsed with distilled water before immersion in each solution. Finally, tissue sections were examined using a light microscope (Nikon), photographed and images were saved as jpg files. For malignant transformation, we applied a score ranging from 0 to 2, where score 0 indicated no regenerated/malignant cell (Fig. 2A); score 1 indicated the presence of liver regenerated nodules (Fig. 2B) and score 2 denoted definitive liver malignancy (Fig. 2C).

**Fig. 2.**
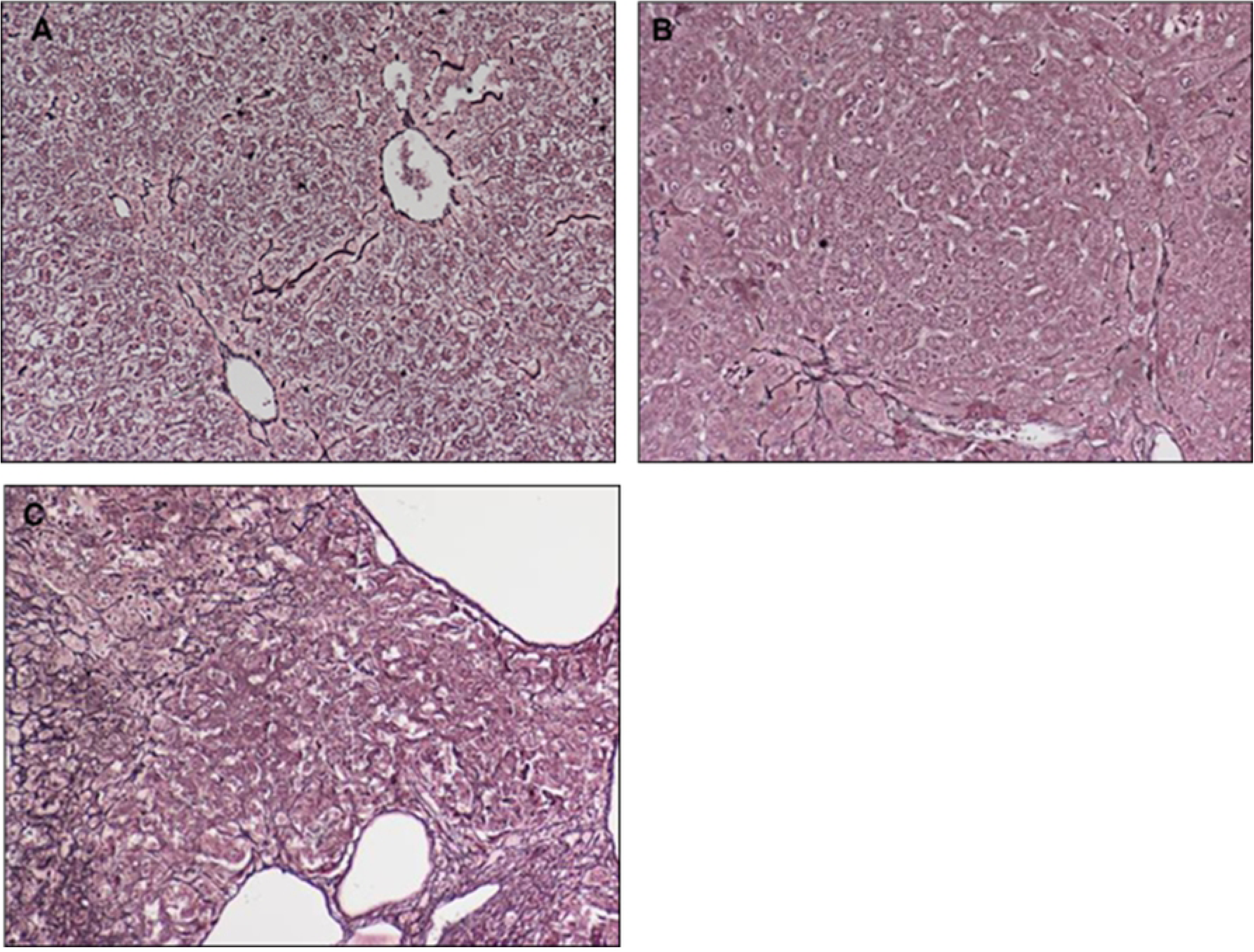
The reticulin staining in hamster liver: We applied a score ranging from 0 to 2, where score 0 indicated no regenerated/malignant cell (Fig. 2A); score 1 indicated the presence of liver regenerated nodules (Fig. 2B) and score 2 denoted definitive liver malignancy (Fig. 2C).

### Immunohistochemistry

We obtained 3 μm tissue sections from paraffin-embedded samples that were subjected to deparaffinisation and rehydration. Following antigen retrieval by autoclaving for 5 min in DAKO antigen retrieval solution (DAKO, Carpenteria, CA) the sections were washed twice in TBS buffer. Endogenous peroxidase was blocked by immersing the slides in 3% hydrogen peroxide solution for 5 min. After washing with TBS, the slides were incubated with primary antibody against OV-6 (mouse anti-OV-6; MAB2020; R&D; 1:200) for 1 hr at room temperature and subsequently washed two times with TBS and incubated with biotinylated secondary antibody for 30 min. After washing two times with TBS, sections were treated with DAB for 5 min. Immediately after staining, the sections were counterstained with haematoxylin for 90 seconds, immersed in xylene and mounted using permount (Fisher Scientific, Pittsburg, PA). Sections were examined using a light microscope (Nikon), photographed and images were saved as jpg files. OV-6-immunostained tissue sections were scored as follows: Score 0 indicated no positive cells (Fig. 3A); score 1 indicated ‘ ≤ 5 stem cells in one high power field’ (Fig. 3B); score 2 indicated ‘ ≥5 and <10 stems cells in one power field’ (Fig. 3C); score 3 indicated ‘>10 stem cells in one power field’ (Fig. 3D). Scores 0 and 1 were considered low-level expression, whereas scores 3 and 4 were considered high-level expression.

**Fig. 3.**
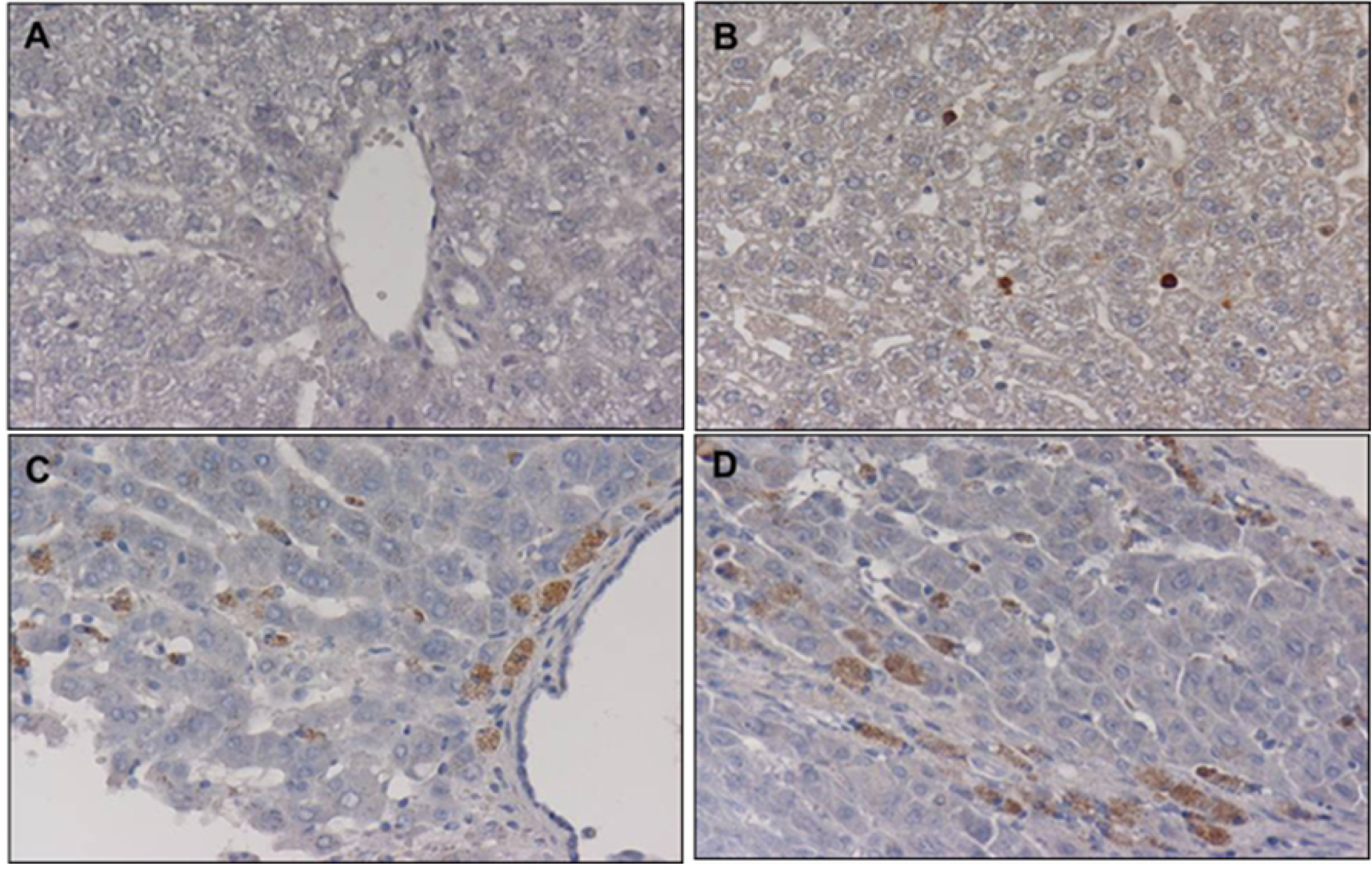
The immunohistochemical staining of OV-6 in hamster liver: Score 0 indicated no positive cells (Fig. 3A); score 1 indicated ‘ ≤ 5 stem cells in one high power field’ (Fig. 3B); score 2 indicated ‘ ≥5 and <10 stems cells in one power field’ (Fig. 3C); score 3 indicated ‘>10 stem cells in one power field’ (Fig. 3D). Scores 0 and 1 were considered low-level expression, whereas scores 3 and 4 were considered high-level expression.

**Fig. 4.**
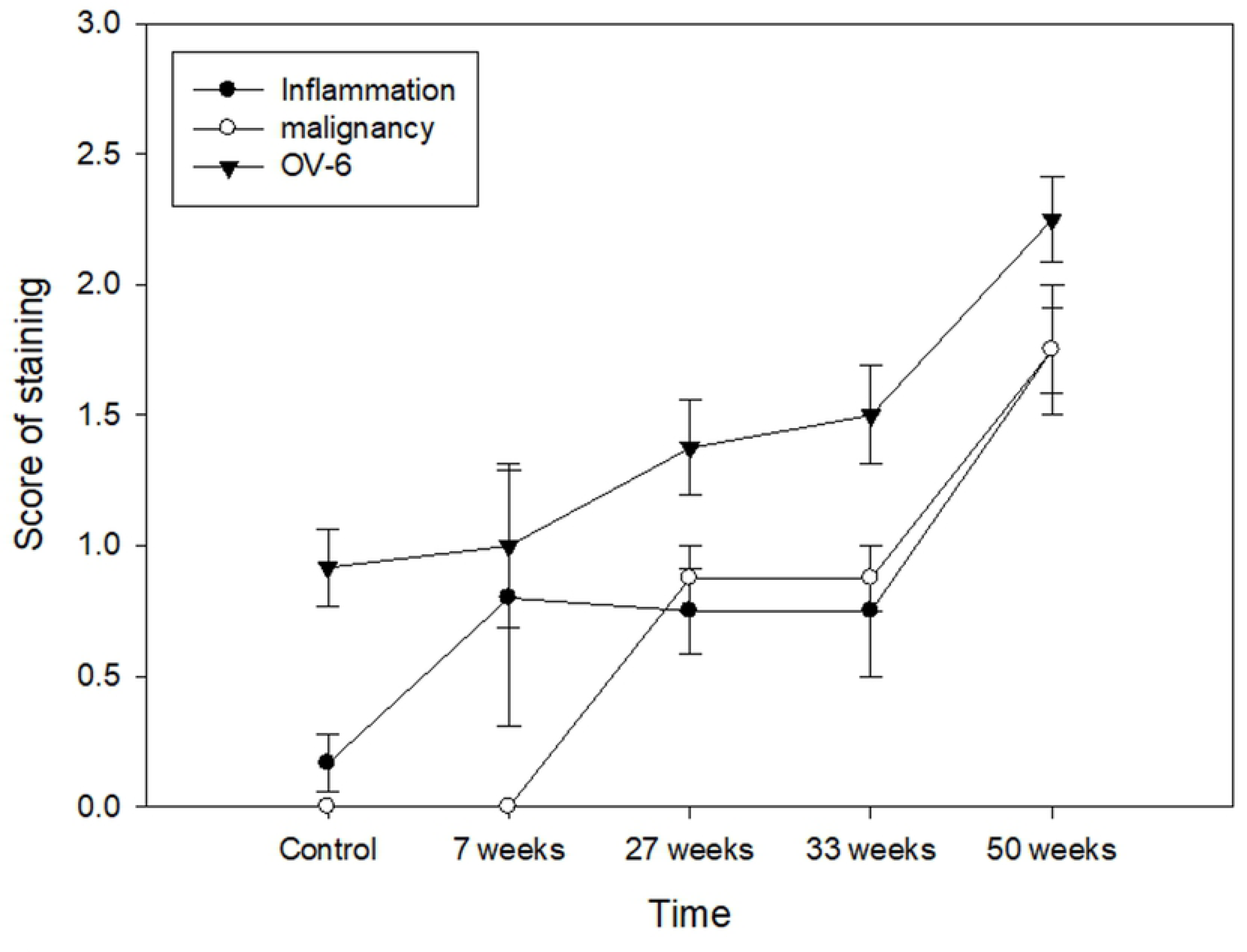
Correlation between malignant transformation, cancer stem cells and inflammation. *P* value was calculated by Spearman’s correlation test. *P* < 0.05 is statistically significant.

### Data Analyses

The relationships between malignant transformation, inflammation and OV-6 were determined using Spearman’s correlation test. Statistical significance was considered when *P* < 0.05.

## Results

### DEN-induced inflammation in liver

Inflammation was determined by examining the scores obtained in H&E-stained sections. We found that DEN progressively induced inflammation at 7 week 7 (40%, 2/5), week 27 (75%, 6/8), week 33 (62.5%, 5/8) and week 50 (100%, 12/12).

### DEN induces malignant transformation in the liver

Malignant transformation was determined by examining the scores obtained from reticulin-stained sections. We found that DEN progressively induced malignant transformation at week 7 (0%, 0/5), week 27 (87.5%, 7/8), week 33 (100%, 8/8) and week 50 (100%, 12/12) (Fig. 2D).

### DEN enhances cancer stem cell marker in the liver

Cancer stem cells were identified based on the immunohistochemical expression of the biomarker, OV-6. Data obtained showed that DEN progressively induced high-levels of OV-6 expression at week 7 (20%, 1/5), week 27 (37.5%, 3/8), week 33 (50%, 4/8) and week 50 (100%, 12/12).

### Relationship between cancer stem cell, inflammation and malignant transformation

DEN-induced inflammation, malignant transformation and high-level OV-6 expression in hamster liver as shown above and applying Spearman’s correlation to the data showed that expression of OV-6 was significantly correlated to inflammation (*p* = 0.001) and malignant transformation (*p* < 0.001)

## Discussion

The goal of this study was to determine the relationship between malignant transformation, inflammation and the role of cancer stem cells in the pathogenesis of HCC. In order to do so, we used DEN as a carcinogen in an experimental hamster model to induce liver tumour, and in this study, we showed that expression of expression of OV-6 was significantly correlated to inflammation and malignant transformation in the DEN-induced hamster model of carcinogenesis.

Liver cancer is the third leading cause of cancer death and the fifth most common cancer worldwide^15^. Because the mechanisms of pathogenesis are not clearly known, the therapeutic approaches for hepatic malignancies are limited. To elucidate this further, the possible mechanisms inducing human HCC are classified into four groups: growth factors and their receptors (TGF-α)^16, 17^, reactivation of developmental pathways (Wnt)^18^, oncogenes (K-*ras*)^19^ and tumour suppressor gene (p53)^20, 21^.

Cancer stem cells are able to self-renew, proliferate, metastasise, cause relapse and induce resistance to chemotherapy and radiation therapy^22^. Cancer stem cells have been identified in various human cancers, including that of the breast^23^, colon^24^, prostate^25^, pancreas^26^, as well as in head and neck squamous cell carcinoma^27^. We showed in this study that DNE-induced HCC expressed more OV-6 positive cells than in control animals, indicating the increased presence of cancer stem cells in DEN-induced model of HCC.

Inflammation is associated with increased risk for various malignant neoplasms, carcinogenesis, metastasis, angiogenesis, tumour invasion, anti-apoptosis, epigenetic modifications, genomic instability, enhanced cell proliferation and aggressive tumour neovascularisation^28–30^. Immune cells including lymphocytes and macrophages, platelets, fibroblasts and tumour cells are a major source of angiogenic factors^31–33^, which play an essential role in leukocyte infiltration into the tumour microenvironment, thereby regulating the tumour size, distribution and composition. Indeed, tumour-associated macrophages are key regulators of the link between inflammation and carcinogenesis^34, 35^.

The activation of hepatic progenitor cells in chronic hepatitis C infection is a common occurrence that depends on the hepatitis stage^36^. Stem cell numbers increase with sinusoidal or interstitial inflammation, and particularly in HCC, are located within liver tumours and scattered as single cells, and not in the portal tracts, bile ducts or canals of Hering. These stem cells assume the morphology of their neighbouring hepatocytes in both cirrhosis and HCCs^37^ and proliferate in fibrous areas and liver tumour parenchyma as the tumours developed. In the current study, reactive ductules and intermediate hepatocyte-like cells originate partly from activation and differentiation of ‘progenitor cells’ in the hamster liver, whose proliferation is associated with an increase in inflammation or the damage of hepatocytes and the tumour status.

In conclusion, there was a significant correlation between number of cancer stem cells, inflammation and malignant transformation in a DEN-induced model of hepatic carcinogenesis in the hamster. It is postulated that cancer developed through an inflammatory process, which increased cancer stem cells within liver tumours. This study describes a simple method for inducing liver tumours, with the objective of identifying and characterising the possible mechanism of malignant transformation.

## Ethics approval and consent to participate

Not applicable.

## Consent for publication

Not applicable.

## Acknowledgements

Not applicable.

